# RNAi Gene knockdown of R-opsin and F-actin in the marine annelid *Streblospio benedicti* by delivering dsRNA

**DOI:** 10.1101/2025.09.22.677873

**Authors:** Jose Maria Aguilar-Camacho, Christina Zakas

## Abstract

**Background:** RNA interference (RNAi) is a genetic tool to disrupt the expression of selected genes by delivering dsRNA into a specific tissue of an organism, resulting in a gene knockdown. We apply RNAi methods to the marine polychaete S*treblospio benedicti*, which is a model system for studying evolutionary and developmental biology, which currently has no established methods for gene expression manipulation.

**Results:** Here we describe a RNAi gene knockdown methodology using two different approaches depending on developmental stage. We feed bacteria expressing dsRNA to early swimming larvae and microinject *in vitro* transcribed dsRNA in juveniles. We used two genes for testing: R-opsin and F-actin. For both developmental stages, gene knockdown was validated using qPCR (quantitative Real-Time PCR). For larvae, we also visualized reduction in RNA expression using Hybridization Chain Reaction (HCR) *in situ* hybridization.

**Conclusions:** We show that both larval feeding and juvenile microinjections are sufficient delivery mechanisms for dsRNA to result in a RNAi gene knockdown. The efficiency can vary depending on the gene and timing, but this is typical for RNAi based approaches.

## 1. INTRODUCTION

RNA interference (RNAi) is a well-studied cytological resistance response to double-stranded RNA (dsRNA) that has been widely used for targeted gene expression knock downs. RNAi is a genetic tool to disrupt the expression of selected genes, and the RNAi pathway is conserved in distinct eukaryotic organisms (Mocellin and Provenzano, 2004; Shan et al. 2010; Lebenzon and Toxopeus, 2024). Due to the efficiency of this transgenic manipulation, many methods have been developed to introduce dsRNA: it can be synthetized *in vitro* and delivered into a specific tissue (i.e., embryos or coelomic cavity) by soaking, injecting, spraying, or feeding (Timmons et al. 2001; Kim et al. 2005; Lebenzon and Toxopeus, 2024). However, some taxonomic groups, such as the Lophotrochozoa, have only a few reported successes with these approaches (Feng et al. 2019).

Feeding dsRNA-expressing bacteria to animals is an effective delivery method in many invertebrate species including insects, nematodes, planarians, and others (Grishok and Mello, 2002; Newmark et al. 2003; Rivera et al. 2011; Iryani et al. 2017; Feng et al. 2019). In aquatic animals, gene knockdowns through bacterial feeding have been successful, such as in the freshwater microcrustacean *Daphnia melanica* (Schumpert et al. 2015), the planarian *Schmidtea mediterranea* (Newmark et al. 2003), the rotifer *Brachionus plicatilis* (Zhang et al. 2024); and specifically for filter-feeding invertebrates such as the oyster *Crassostrea gigas* (Feng et al. 2019), the marine sponge *Tethya wilhema*, and the freshwater sponge *Ephydatia* fluviatilis (Rivera et al. 2011). The methods for feeding dsRNA-expressing bacteria in these species varies widely across experiments, but in all cases gene expression was successfully knocked down (Feng et al. 2019).

The marine annelid *Streblospio benedicti* is a model system for studying developmental evolution and evolutionary genetics, but to date, no methods for gene expression manipulation have been established. This annelid has two distinctively different offspring morphs that differ dramatically in their ontogeny, life-history, and morphology (reviewed in Zakas 2022). The planktotrophic larval morph is an obligately feeding and swimming larva that develops over weeks in the plankton, while the lecithotrophic larval morph is facultatively (non-obligately) feeding with an abbreviated larval phase and notable morphological and behavioral differences (Zakas et al. 2018). As a model for developmental evolution, *S. benedicti* is easily cultured in the lab (McHugh, 2024); although one drawback for establishing functional genetic tools is that the timing of fertilization cannot be controlled in the lab, so typical methods of early embryo microinjections are problematic (Zakas, 2022). For species where microinjections are difficult, or later developmental stages are necessary, delivery of dsRNA through other means becomes more efficient. Here we present two methods of delivery of dsRNA to knock down gene expression in *S. benedicti*.

We established a gene knockdown by RNAi methodology in *S. benedicti* using two methods depending on developmental stage. We use the bacteria feeding approach with planktotrophic feeding larvae and microinjection of *in vitro* transcribed dsRNA in juveniles. To test our approaches, we used two genes: R-opsin (expressed in eyespots and involved in light sensing pathways) and F-actin (ubiquitously expressed in the cellular cytoskeleton to transport molecules). For both developmental stages, we validated RNA knock-down efficiency by qPCR. For larvae, we visualized reduction in RNA expression using Hybridization Chain Reaction (HCR*) in situ* hybridization (Choi et al. 2018; Aguilar-Camacho et al. 2024). Our study constitutes the first successful implementation of gene knockdown by RNAi in a marine annelid species, by feeding dsRNA expressing bacteria in free-swimming larvae and microinjecting synthesized dsRNA in juveniles.

## 2. RESULTS AND DISCUSSION

### 2.1 Larval Feeding Approach

We selected the candidate genes R-opsin and F-actin for RNAi trials as these genes have known expression patterns in *S. benedicti*. We selected R-opsin for a candidate gene approach as R-opsin spots can be easily visualized in the head of larval and juvenile animals using HCRs. In *Capitella teleta*, CRISPR-Cas9 injections reduced the expression of the R-opsin spots and the deletion of one of the two eye spots in that species (Neal et al. 2019). F-actin was selected as it is a ubiquitously expressed gene, suitable for RNAi experiments (Rivera et al. 2011). Expression of dsRNA for each gene was induced in *E. coli* strain HT115 (DE3) with 0.4 mM IPTG for 4 h at 37 °C. Total RNA was isolated from induced bacteria and observed via 1% agarose gel electrophoresis (Fig. 1A). A band of dsRNA is visible for each gene indicating the bacteria are expressing dsRNA of the correct size (Fig. 1A). Planktotrophic larvae of *S. benedicti* are obligately-feeding larvae that non-selectively consume single-celled algae (McHugh, 2024). Under typical lab conditions, planktotrophic larvae are fed an abundance of *Tetraselmis* sp. algae until they metamorphose into juveniles approximately two weeks later (McHugh, 2024). We hypothesized that larvae would also ingest suspended bacteria that are expressing dsRNA. Therefore, we used two treatments: larval feeding of bacteria with and without *Tetraselmis* sp. algae included. We sampled two timepoints (4 hours and 12 hours post-feeding).

**Figure 1.**
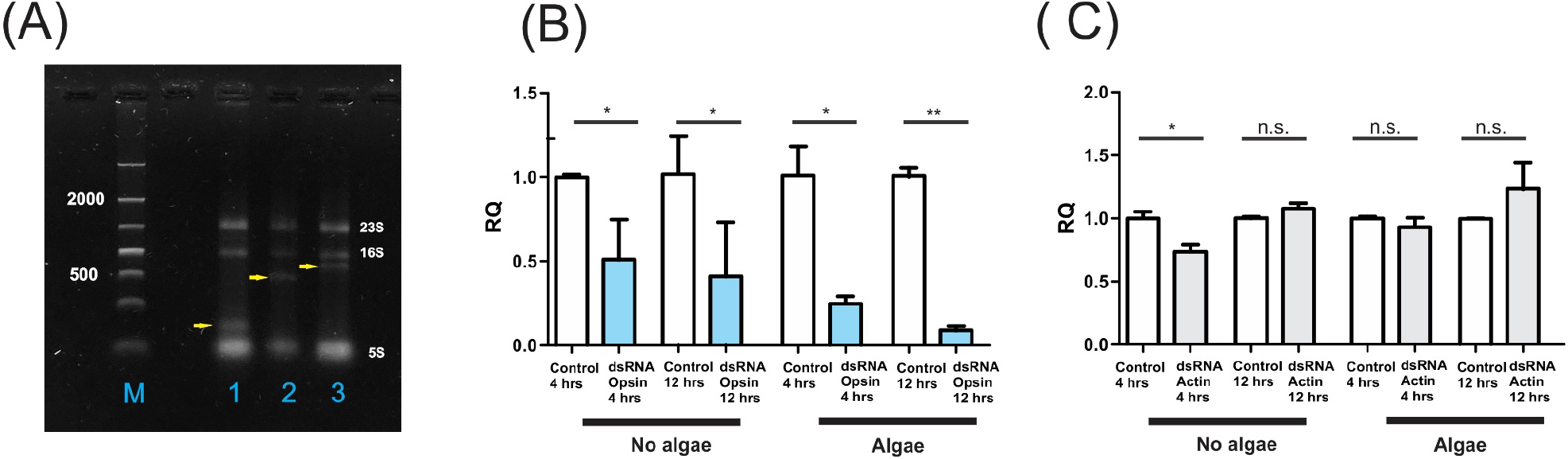
dsRNA Induction and larval feeding knockdown. A) 1% agarose gel loaded with total RNA of induced bacteria. M= 3000bp DNA ladder,1= Control sample (L4440 plasmid with no gene insert), 2= L4440 plasmid with a fragment of the R-opsin and 3 = L4440 plasmid with a fragment of the F-actin. The three bands in lanes 1, 2 and 3 correspond to the ribosomal RNA subunits of the bacteria (23S, 16S and 5S). Yellow arrows indicate the dsRNA expressed for each sample 4 hours after the induction with 0.4 mM IPTG. B) qPCR analysis of the R-opsin expression levels with and without algae at two timepoints post feeding. C) qPCR analysis of the F-actin expression levels with and without algae at two timepoints post feeding Y-axis denotes relative levels of expression (RQ). Bars represent mean ± SE of n= 2 or 3 replicates. Unpaired t-tests (two-tails) were calculated and statistical significances in all cases were defined as: n.s.= no significant differences, *=*P* < 0.05, **= *P* < 0.001.

We fed ‘early planktotrophic’ larvae (3-eye stage larvae that are approximately five days old) bacteria expressing either dsRNA for R-opsin or F-actin for three hours in two separate experiments. As early larvae are capable of swimming and feeding, but have not yet developed a 4^th^ eyespot, this was an ideal stage to test dsRNA knock-down efficiency using R-opsin. Eye spot formation in *S. benedicti* is not fully understood, however the swimming planktotrophic larvae have four visible, darkly pigmented eyespots. The first pair of eyes appear at the trochophore stage, followed by a 3^rd^ eye on the right side, and a 4^th^ on the left approximately one day later (see Aguilar-Camacho et al. 2024). R-opsin patterning seems to overlap with the physical eyespots but with an additional, more posterior, spot on each side (three pairs of spots). It is likely that these spots are in the sub-ectodermal region of the head and are colocalized with the pigmented eyespots for two of the three pairs (Fig. 2A). Similarly, six R-opsin spots (three pairs) are also expressed at the early juvenile stage (Fig. 2B). R-opsin spots are colocalized with the eye spots in the larval stage of other polychaeta species such as: *Platynereis dumerilii* and *C. teleta* (Randel et al. 2013; Neal et al. 2019); but they can also be found in other regions as well (Backfish et al. 2013; Neal et al. 2019).

**Figure 2.**
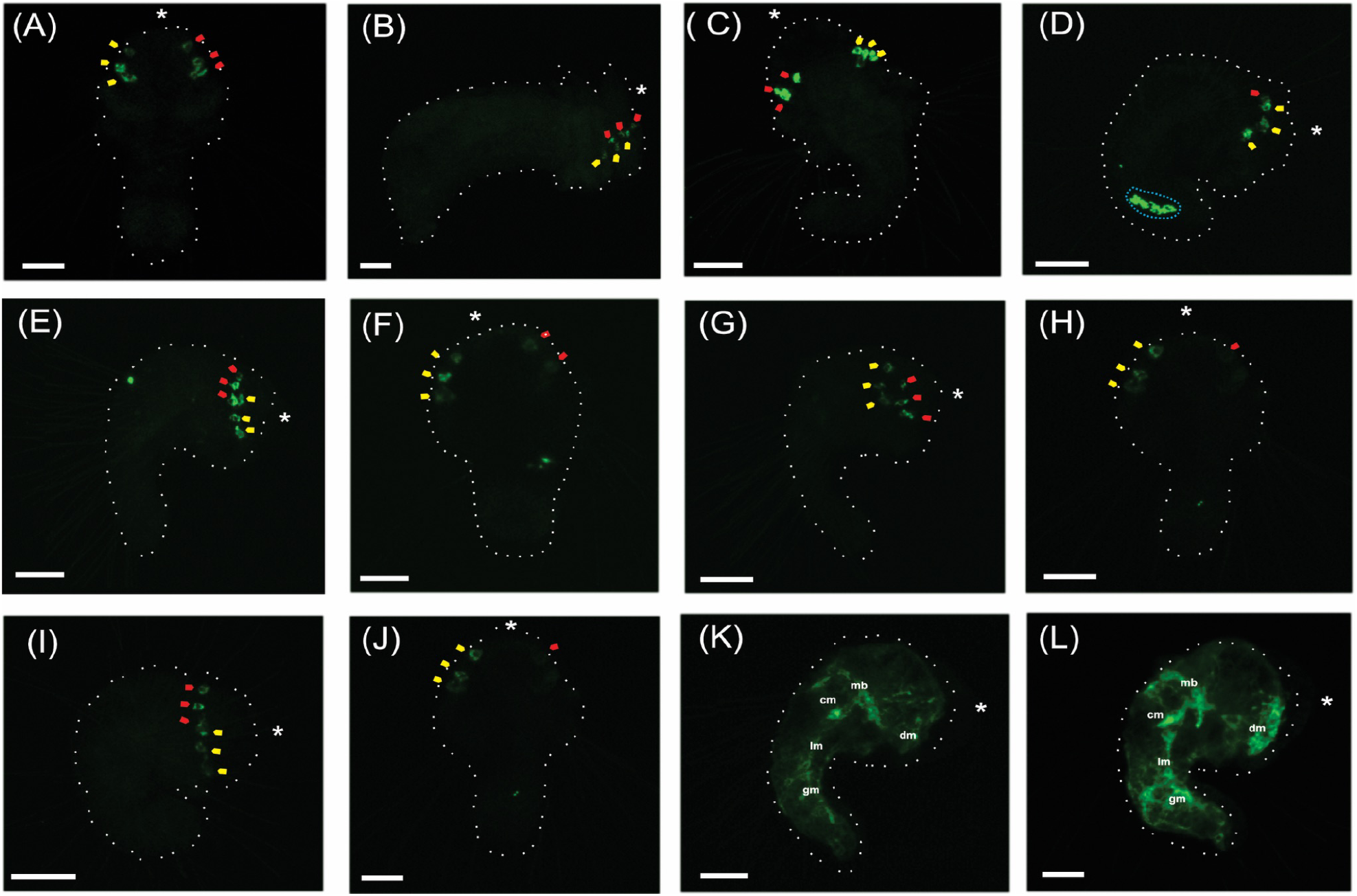
HCR *in situ* hybridization of the *S. benedicti* R-opsin and F-actin from larval feeding. Yellow arrowheads indicate R-opsin spots from the right side and red arrowheads indicate R-opsin spots from the left side. White asterisk indicates the anterior end of the larva. White dashed lines outline the body for each individual sample. Scale bars = 45 µm. A) Wild type ventral view of an early planktotrophic larva showing six R-opsin spots—three per each side. B) Wild type lateral view of a planktotrophic juvenile showing six R-opsin spots. C) Control early planktotrophic larva from the feeding experiment no algae/4h experiment. Dorsal view shows all six R-opsin spots. D) Experimental larva from the R-opsin dsRNA /no algae/4h experiment. Lateral view shows four R-opsin spots— three on the right and one on the left. Blue dotted circle shows consumed bacteria expressing R-opsin dsRNA in the larval gut that reacted with the R-opsin probes. E) Control larva from the no algae/12h experiment. Lateral view shows all six R-opsin spots. F) Experimental larva from the R-opsin /no algae/12h experiment ventral view shows five R-opsin spots— three on the right and one on the left. G) Control larva from the R-opsin/ algae/4h experiment. Lateral view shows six R-opsin spots. H) Experimental larva from the R-opsin dsRNA/algae/4h experiment. Ventral view shows four R-opsin spots— three on the right and one on the left. I) Control larva from the algae/12h experiment. Lateral view shows six R-opsin spots. J) Experimental larva from the R-opsin dsRNA /algae/12h experiment. Ventral view shows four R-opsin spots: three on the right and one on the left. K) Control larva from the F-actin dsRNA/no algae/4h experiment. Lateral view of an early-planktotrophic larva. L) Experimental larva from the F-actin dsRNA/no algae/4h experiment. Lateral view of an early-planktotrophic larva. *lm*=longitudinal muscle, *gm*=gastric muscle, *mb*= muscle band of the head, *cm*=chaetal muscle and *dm*=dense muscle mass of the ventral head.

### 2.2 Larval Feeding Results

We quantified gene expression differences between our treatments using qPCR and HCR *in situ* for larval feeding experiments. Early planktotrophic larvae of the feeding experiment showed a reduction of the expression levels of R-opsin, compared to their respective control samples (unpaired t-test, p<0.05) in all four cases tested based on qPCR (Fig. 1B). In contrast, ‘early planktotrophic’ larvae showed a reduction of the expression levels of F-actin in only one case: 4h after washing the samples that were fed exclusively with bacteria expressing F-actin dsRNA (unpaired t-test, p<0.05) based on qPCR (Fig. 1C). Therefore, three hours of ingesting dsRNA-expressing bacteria in the early planktotrophic larvae was sufficient time for the larvae to ingest bacteria to reduce gene expression (Fig. 1B, C; Supplement Data 1 and 2).

Using HCR *in situs*, we visualized the pattern of RNA expression in the larval feeding approach. We visualized fixed animals at the ‘early planktotrophic’ stage (3-eyed larvae that are approximately five days old, when all six R-opsin spots and three eyes are present). Animals fed bacteria expressing R-opsin dsRNA (with and without algae) showed a reduction in R-opsin spots: only 4-5 viable spots out of six (Fig. 2 D, F, H, J). Interestingly, the reduction of the R-opsin spots is on the left side of the larval head, which is the side that forms the 4^th^ eye later in development. In the treatments, the R-opsin spots on the left side are not as bright as those from the right side (Fig. 2 F, H, J). There is also a noticeable reduction in brightness in the R-opsin spot that does not overlap with an eyespot (Fig. 2 H, J).

F-actin is ubiquitously expressed in cells of early planktotrophic larvae, and it is likely that the expression is in muscle cells of the endodermal region of the larva. There are some anastomosed, irregular bands composed of longitudinal muscles running dorso-laterally towards the posterior end. A muscle band surrounding the head converges with the longitudinal muscle. Chaetal muscles, gastric muscles, and a dense muscle mass in the ventral region of the head are also noticeable (Fig. 2K). After larvae were fed F-actin dsRNA expressing bacteria, no visible changes (based on HCRs) were detected although there were significant differences of the gene expression based on qPCR (Fig. 2K, J). Similar results for actin knockdowns were also reported in sponge and cnidarian species, which showed a reduction of the expression levels of actin based on qPCR, but no noticeable phenotypic differences (based on phalloidin staining) (Rivera et al. 2011 Kyslik et al. 2024). In contrast, a gene knockdown of actin by transfection in the sea anemone *Aiptasia pallida* showed altered tissue morphology in samples with high doses and long incubation time of dsRNA (Dunn et al. 2007). It is possible that a visual actin knockdown phenotype could be achieved in *S. benedicti* with longer exposure or higher dosage of dsRNA than was tested in this experiment.

The different results between the two genes tested here could indicate that ubiquitously expressed or critical genes— which may have genetic redundancy— are less susceptible to knockdown by dsRNA using these conditions. Variability in the efficiency of dsRNA knockdowns between genes within a species is a common phenomenon (Kamath et al. 2000; Lebenzon and Toxopeus, 2024).

### 2.3 Juvenile microinjection approach

We injected *in vitro* transcribed dsRNA (120 ng/ µL) of R-opsin and F-actin into 3-week-old juveniles, as adult tissue injection is another common technique which has been successful in annelids (Chen et al. 2020). We tested gene expression differences using qPCR at two timepoints (6 h and 12 h after microinjection). At both timepoints the R-opsin injected animals showed a reduction of the expression levels (unpaired t-test: P<0.05; Fig. 3A). Animals injected with F-actin dsRNA also had reduced expression (unpaired t-test: P<0.05) but only at one time point, 12h after injection (Fig. 3B).

**Figure 3.**
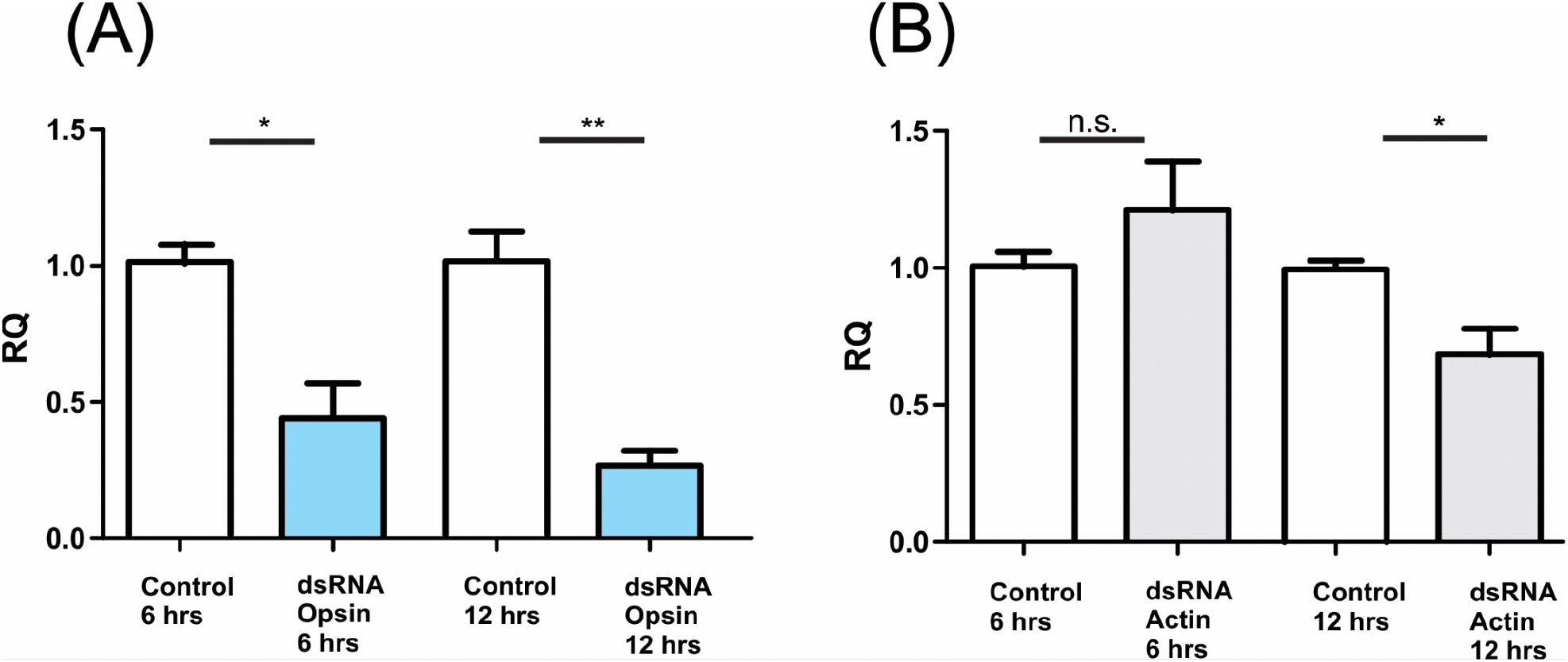
qPCR Expression levels for microinjected juveniles. A) qPCR analysis of the R-opsin expression levels. B) qPCR analysis of the F-action expression levels. Y-axis denotes relative levels of expression (RQ). Bars represent mean ± SE of n= 3 replicates. Unpaired t-tests (two-tails) were calculated and statistical significances in all cases were defined as: n.s.= no significant differences, *=*P* < 0.05, **= *P* < 0.001.

A gene knockdown of R-opsin was observed at the two timepoints after microinjection of dsRNA, and a gene knockdown of F-actin only at the second timepoint (Figs. 3A, B; Supplement Data 3 and 4). These results again demonstrate that there is variability in the knockdown efficiency and timing between genes, but the delivery method by microinjection is sufficient to achieve expression reduction in the target gene. In selected species of arthropods, differences in the knockdown efficiency of opsin and actin genes are dependent on the dsRNA concentration and time after injection (Fan et al. 2022; Huang et al. 2023; Abba et al. 2019; Rosa et al. 2012).

## 3. CONCLUSIONS

We have demonstrated that both larval feeding and juvenile microinjection are sufficient delivery mechanisms for dsRNA to result in RNAi in target genes. This is a major step for transgenic experiments in post-embryonic developmental stages in marine species. The efficiency of gene knockdown can vary depending on the gene, timing, or dosage of dsRNA, but this is typical for RNAi based approaches. We can conclude that an appropriate delivery method depends on the target gene and the developmental stage but dsRNA for RNAi knock-down is achievable in *S. benedicti*.

## 4. EXPERIMENTAL PROCEDURES

### 4.1 Animal culturing

Early planktotrophic larvae and 3-week-old juveniles (planktotrophic morph) are selected from maintained laboratory cultures.

### 4.2 Feeding early planktotrophic larvae with bacteria expressing dsRNA

R-opsin (1224 bp) and F-actin (1131 bp) genes were identified in the transcriptome of *S. benedicti* (as in Aguilar-Camacho et al. 2024; Supplement Data 7). A section R-opsin (467 bp) and F-actin (584 bp) were synthesized (Genscript, USA) and inserted into the L4440 (Abgene, USA) plasmid vector (SacI-HindIII). This vector contains two convergent T7 polymerase promoters in opposite orientation separated by a multicloning site which is used for the induction of dsRNA in *E. coli* cells (Timmons et al. 2001). HT115 (DE3) competent cells were prepared using standard CaCl_2_ methodology and were transformed with the plasmid vectors by heat shock. Transformed cells were grown overnight onto LB agar plate with Ampicillin (100 μg/mL) at 37°C. A positive bacteria colony for each gene was chosen for an overnight culture in a 5 mL LB medium with Ampicillin (100 μg/mL) and Tetracycline (50 μg/mL) at 37°C. The next day, the overnight bacteria culture was inoculated (500 μL) into a 50 mL LB medium with Ampicillin (100 μg/mL) and Tetracycline (50 μg/mL) at 37°C until OD_595_= 0.4 was reached. Expression of dsRNA was induced with 0.4 mM IPTG for 4 h at 37°C under agitation. Induced bacteria cultures were centrifuged (3000 rpm × 3 minutes) and the pellet washed twice with artificial sea water.

For larval feeding, ∼1 × 10^8^ cells/mL of *E. coli* were added to larval cultures with and without *Tetraselmis* sp. algae (∼2000 cells/mL; Carolina Biology, US) for 3 h in a well from a six-well plate (planktotrophic larvae of *S. benedicti* are fed *Tetraselmis* sp. algae in lab culture). The duration was chosen based on preliminary experiments in which bacteria expressing dsRNA were added to planktotrophic larvae in a well, and after 3 h some of the larvae died and a dense layer (probably bacteria) was precipitated at the bottom of the well. After feeding planktotrophic larvae for 3 h, the larvae were washed twice with artificial seawater. Gene expression was measured at two timepoints, 4 h and 12 h after washing the larvae samples that were fed; two treatments (with and without algae) and two to three replicates for each timepoint and treatment. Control samples were fed with induced bacteria from a transformed positive colony with the L4440 plasmid where no gene was inserted. Clutches of planktotrophic larvae can be >250 individuals, which were divided into treatments and controls of 25 individuals per well. At the end of the experiment, some larvae were fixed for HCR imaging, and others were collected for RNA extraction for qPCR.

### 4.3 Microinjection of *in vitro* synthesized dsRNA into juveniles

The R-opsin (467 bp) and F-actin (602 bp) fragment was amplified by PCR from cDNA of a 3-week-old juvenile using the following primers: 5’ TAATACGACTCACTATAGGGCCAGTACCAAGCACCTTCGGAC (forward for R-opsin) and 5’ TAATACGACTCACTATAGGGGAAGAACATAGGCACTCCAAAC (reverse for R-opsin), 5’ TAATACGACTCACTATAGGGGACGATGATGTTGCTGCTCTCG (forward for F-actin), 5’ TAATACGACTCACTATAGGGCGCTCGGTAAGGATCTTCATCA (reverse for F-actin). PCR products for each gene were purified by using the DNA Clean & Concentrator kit (Zymo Research, USA) and used as template for dsRNA in vitro transcription using the HiScribe® T7 High Yield RNA Synthesis Kit (NEB, USA). The RNA pellet for each gene was resuspended in 50 μL of Molecular Biology Water (Sigma, USA) and stored at −80°C until further use. We chose 3-week-old juveniles for injection as other developmental stages, such as early juveniles and swimming lecithotrophic larvae leaked the solution after microinjection. In contrast, 2-month-old juveniles and adults have a dense epidermal cuticle that is difficult to penetrate. Prior to injection, 3-week-old juveniles were relaxed by using 700 mM of MgCl_2_ (Sigma, USA) with artificial sea water for 30 minutes at room temperature. Each worm was then placed into a small petri dish with artificial sea water in which half of the petri dish was filled with 3% Molecular Biology grade agarose (Apex, US). This agarose structure was used as a “wall” and helped throughout the injection of the worms. Microinjection was applied with an InjectMan® 4 Micromanipulation Workstation (Eppendorf, Germany) connected to a Femtojet® 4i system (Eppendorf, Germany) that was observed using an AXIO Observer A1 Inverted Fluorescence Microscope (Zeiss, Germany). Heat pulled quartz needles (Sutter Instrument, USA) were backfilled with 120 ng/ µL dsRNA of the target gene (either R-opsin or F-actin) and 1.5% (W/V) Red Rhodamine-Dextran (70 000MW, Neutral, Invitrogen, US). Control samples were only injected with Rhodamine-Dextran. The solution was injected (pi 150, ti 1.20, pc 60) in the gastric cavity of each worm as well as in two segments that were located three to five segments below the prostomium, and a single individual worm was considered as one replicate (Fig. S1 A, B). We tested gene expression differences at two timepoints (6 h and 12 h after injection) with three biological replicates for the control and experiment samples for each gene tested. After injection, individuals were transferred into a small petri dish with artificial seawater and stored in a small incubator at 20°C until they were used for total RNA extraction.

### 4.4 Total RNA extraction and qPCR analyses

For all experiments, total RNA was extracted using the Monarch ® Total RNA prep kit (NEB, USA). To test for dsRNA expression, induced bacteria (<5 × 10^7^) were centrifuged at 3000 rpm × 3 minutes in an Eppendorf tube and a portion of the supernatant removed for RNA extraction. Each replicate of the feeding experiment and the injected juvenile worms were transferred into an Eppendorf tube and centrifuged at 2000 rpm × 3 minutes. Artificial seawater was removed prior to RNA extraction. RNA was extracted following the Monarch® (NEB, US) protocol. RNA was stored at −80°C until further use. Total RNA from bacteria expressing dsRNA was loaded into a 1% Molecular Biology grade agarose gel (Apex, US) for 45 min at 80V and the gel imaged using a Accuris Instruments SmartDoc 2.0 (Nova Tech, USA). Reverse transcription (RT) was carried out using the Monarch® Protoscript II First Strand cDNA synthesis Kit (NEB, USA). RNA quantifications were determined by using the Qbit 4 Fluorometer (Thermo Fisher, USA). Primers for R-opsin are: 5’ CCTCACGGTCAACAGCATCT (forward) and 5 ‘CTGCCTTCATGGGGTTGGAT (reverse), 158 bp product length. Primers for F-actin are: 5’ TGGTGGGTATGGGCCAGAAG (forward) and 5’ CCAACTGGGACGACATGGAGA (reverse), 120 bp product length. Three independent repeats for each replicate were examined and the samples were subjected to 40 cycles of amplification in a QuantStudio 6 Pro Real-Time PCR System (Thermo Fisher, USA) and the Kapa SYBR Fast qPCR master mix was used (Roche, Switzerland). All relative levels of gene expression of the R-opsin and F-actin were normalized to the housekeeping gene Elongation Factor 1 (EF-1) with the control samples for each independent experiment (Ct values of EF-1 were stable in all the samples tested). Primers of EF-1 are: 5 ‘TGTGGAGACTTTCAGCGAGT (forward) and 5’ CGAGGCTCACTTCTTCTTGC (reverse), 163 bp product length. Data were analyzed using the ΔΔCt method and the relative levels of expression were calculated by the 2^−ΔΔCt^ method, which is expressed as RQ (relative quantity) (Rao et al. 2013; Livak and Schmittgen, 2001).

### 4.5 Statistical analyses

RQ data values (2^−ΔΔCt^ or 2^−ΔΔCt-1^) were compared between the samples of the experiment with their respective RQ data values of the control samples. Unpaired t-tests (two-tails) were calculated and statistical significances in all cases were defined at *P* < 0.05. These analyses were performed with the GraphPad Prism software (Supplement Data 1, 2, 3 and 4).

### 4.6 Hybridization chain reaction (HCR) *in situ* hybridization and Imaging

Multiplex probes of R-opsin and F-actin transcripts of *S. benedicti* were designed using the HCR3.0 Probe Maker (Kuehn et al. 2022; Supplement Data 5 and 6). Sequences were synthesized into DNA oligo pools (50 pmol DNA Pools Oligo Pool) from Integrated DNA Technologies and resuspended to 1 pmol/μl in nuclease free water (NEB, USA). Animals were fixed in 4% paraformaldehyde at 4 °C overnight, transferred stepwise into 100% methanol, and kept at − 20 °C. Samples were rehydrated through a methanol/DEPC-treated PBSt series. HCR was performed as previously described (Aguilar-Camacho et al. 2024). HCR samples were mounted in Slowfade Glass (Thermo Fisher, USA) with DAPI and kept at 4 °C until imaging and imaged using Zeiss Laser Scanning Confocal Microscope LSM 880. Z-stack images (40 layers) were processed in ImageJ (Sheffield, 2007). We imaged two-to-five individuals of each sample, for each gene (R-opsin and F-actin), from each treatment and timepoint of the feeding experiment.

## ACKNOWLEDGEMENTS

We thank Kayleigh McHugh, Liv Borges, Robert Walshmith, and Nathan Harry for help with animal collection and maintenance of samples. Caitil Heil and Nathan Brandt assisted with the bacteria culture. Mariusz Zareba at the NC State CMIF, NSF DBI-1624613, assisted with the confocal analysis grant. This work was supported by NIH NIGMS 5R35GM142853 to C. Zakas.

## REFERENCES

Abbà, S., Galetto, L., Ripamonti, M., Rossi, M., Marzachí, C. (2019). RNA interference of muscle actin and ATP synthase beta increases mortality of the phytoplasma vector Euscelidius variegatus. Pest Management Science, 75(5), 1425–1434.

Aguilar-Camacho, J. M., Harry, N. D., Zakas, C. (2024). Comparative Hox genes expression within the dimorphic annelid Streblospio benedicti reveals patterning variation during development. EvoDevo, 15(1), 12.

Backfisch, B., Veedin Rajan, V. B., Fischer, R. M., Lohs, C., Arboleda, E., Tessmar-Raible, K., Raible, F. (2013). Stable transgenesis in the marine annelid Platynereis dumerilii sheds new light on photoreceptor evolution. Proceedings of the National Academy of Sciences, 110(1), 193–198.

Chen, C. P., Fok, S. K. W., Hsieh, Y. W., Chen, C. Y., Hsu, F. M., Chang, Y. H., Chen, J. H. (2020). General characterization of regeneration in Aeolosoma viride (Annelida, Aeolosomatidae). Invertebrate Biology, 139(1), e12277.

Choi, H. M., Schwarzkopf, M., Fornace, M. E., Acharya, A., Artavanis, G., Stegmaier, J., Cunha, A.,Pierce, N. A. (2018). Third-generation in situ hybridization chain reaction: multiplexed, quantitative, sensitive, versatile, robust. Development, 145(12), dev165753.

Dunn, S. R., Phillips, W. S., Green, D. R., Weis, V. M. (2007). Knockdown of actin and caspase gene expression by RNA interference in the symbiotic anemone Aiptasia pallida. The Biological Bulletin, 212(3), 250–258.

Fan, Z., Zhang, Z., Zhang, X., Kong, X., Liu, F., Zhang, S. (2022). Five Visual and Olfactory Target Genes for RNAi in Agrilus planipennis. Frontiers in Genetics, 13, 835324.

Feng, D., Li, Q., Yu, H. (2019). RNA interference by ingested dsRNA-expressing bacteria to study shell biosynthesis and pigmentation in Crassostrea gigas. Marine Biotechnology, 21, 526–536.

Grishok, A., Mello, C. C. (2002). RNAi (nematodes: Caenorhabditis elegans). Advances in genetics, 46, 339–360.

Huang, M., Meng, J. Y., Zhou, L., Yu, C., Zhang, C. Y. (2023). Expression and function of opsin genes associated with phototaxis in Zeugodacus cucurbitae Coquillett (Diptera: Tephritidae). Pest Management Science, 79(11), 4490–4500.

Iryani, M. T. M., MacRae, T. H., Panchakshari, S., Tan, J., Bossier, P., Wahid, M. E. A., Sung, Y. Y. (2017). Knockdown of heat shock protein 70 (Hsp70) by RNAi reduces the tolerance of Artemia franciscana nauplii to heat and bacterial infection. Journal of experimental marine biology and ecology, 487, 106–112.

Kamath, R.S., Martinez-Campos, M., Zipperlen, Fraser, A.G., & Ahringer J. (2000). Effectiveness of specific RNA-mediated interference through ingested double-stranded RNA in Caenorhabditis elegans. Genome Biology, 2, 2000.

Kim, D. H., Behlke, M. A., Rose, S. D., Chang, M. S., Choi, S., Rossi, J. J. (2005). Synthetic dsRNA Dicer substrates enhance RNAi potency and efficacy. Nature biotechnology, 23(2), 222–226.

Kuehn, E., Clausen, D. S., Null, R. W., Metzger, B. M., Willis, A. D., Özpolat, B. D. (2022). Segment number threshold determines juvenile onset of germline cluster expansion in Platynereis dumerilii. Journal of Experimental Zoology Part B: Molecular and Developmental Evolution, 338(4), 225–240.

Kyslík, J., Born-Torrijos, A., Holzer, A. S., Kosakyan, A. (2024). RNAi-directed knockdown in the cnidarian fish blood parasite Sphaerospora molnari. Scientific Reports, 14(1), 3545.

Lebenzon, J. E., Toxopeus, J. (2024). Knock down to level up: Reframing RNAi for invertebrate ecophysiology. Comparative Biochemistry and Physiology Part A: Molecular & Integrative Physiology, 297, 111703.

Livak, K. J., Schmittgen, T. D. (2001). Analysis of relative gene expression data using real-time quantitative PCR and the 2− ΔΔCT method. methods, 25(4), 402–408.

McHugh, Kayleigh. (2024). “Animal Care Protocol: Streblospio benedicti.” dx.doi.org/10.17504/protocols.io.eq2lyw24wvx9/v1

Mocellin, S., Provenzano, M. (2004). RNA interference: learning gene knock-down from cell physiology. Journal of translational medicine, 2, 1–6.

Neal, S., De Jong, D. M., Seaver, E. C. (2019). CRISPR/CAS9 mutagenesis of a single r-opsin gene blocks phototaxis in a marine larva. Proceedings of the Royal Society B, 286(1904), 20182491.

Newmark, P. A., Reddien, P. W., Cebria, F., Alvarado, A. S. (2003). Ingestion of bacterially expressed double-stranded RNA inhibits gene expression in planarians. Proceedings of the National Academy of Sciences, 100(suppl_1), 11861–11865.

Randel, N., Bezares-Calderón, L. A., Gühmann, M., Shahidi, R., Jekely, G. (2013). Expression dynamics and protein localization of rhabdomeric opsins in Platynereis larvae. Integrative and Comparative Biology. 53(1):7–16.

Rao, X., Huang, X., Zhou, Z., Lin, X. (2013). An improvement of the 2^ (–delta delta CT) method for quantitative real-time polymerase chain reaction data analysis. Biostatistics, bioinformatics and biomathematics, 3(3), 71.

Rivera, A. S., Hammel, J. U., Haen, K. M., Danka, E. S., Cieniewicz, B., Winters, I. P., Posfai, D., Wörheide, G., Lavrov, D.V., Knight, S.W., Hill, M.S., Hill, A.L., Nickel, M. (2011). RNA interference in marine and freshwater sponges: actin knockdown in Tethya wilhelma and Ephydatia muelleri by ingested dsRNA expressing bacteria. BMC biotechnology, 11, 1–15.

Rosa, C., Kamita, S. G., Falk, B. W. (2012). RNA interference is induced in the glassy winged sharpshooter Homalodisca vitripennis by actin dsRNA. Pest management science, 68(7), 995–1002.

Schumpert, C. A., Dudycha, J. L., Patel, R. C. (2015). Development of an efficient RNA interference method by feeding for the microcrustacean Daphnia. BMC biotechnology, 15, 1–13.

Shan, G. (2010). RNA interference as a gene knockdown technique. The international journal of biochemistry and cell biology, 42(8), 1243–1251.

Sheffield, J. B. (2007). ImageJ, a useful tool for biological image processing and analysis. Microscopy and Microanalysis, 13(S02), 200–201.

Timmons, L., Court, D. L., Fire, A. (2001). Ingestion of bacterially expressed dsRNAs can produce specific and potent genetic interference in Caenorhabditis elegans. Gene, 263(1-2), 103–112.

Zakas, C., Deutscher, J. M., Kay, A. D., & Rockman, M. V. (2018). Decoupled maternal and zygotic genetic effects shape the evolution of development. Elife, 7, e37143.

Zakas, C. (2022). Streblospio benedicti: A genetic model for understanding the evolution of development and life-history. In Current Topics in Developmental Biology (Vol. 147, pp. 497–521). Academic Press.

Zhang, Y., Kan, D., Zhou, Y., Lian, H., Ge, L., Shen, J., Dai, Z., Shi, Y., Han, C., Yang, J. (2024). Efficient RNA interference method by feeding in Brachionus plicatilis (Rotifera). Biotechnology Letters, 1–11.

