## Supplemental Fig1 for "RNAi Gene knockdown of R-opsin and F-actin in the marine annelid *Streblospio benedicti* by delivering dsRNA"

**Figure Supplement 1**. **Microinjection of dsRNA and dextran into 3-week-old juveniles of *S. benedicti*.** A) Lateral view of an individual injected in the gastric cavity. B) Dorsal view of an individual injected in two body segments located five segments below the prostomium. Green arrows indicate the solution injected that contains dsRNA+dextran. Scale bars= 100 µm.

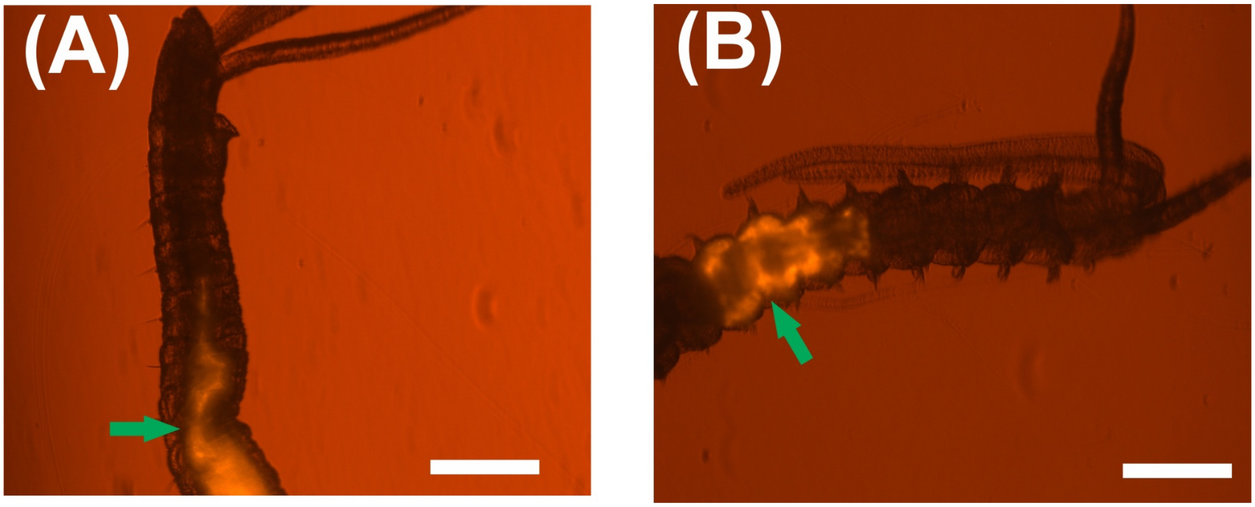
